# Recurrent enhancer-promoter interactions across samples

**DOI:** 10.1101/2025.09.27.678855

**Authors:** Mathew Weston, Satvik Gunjala, Haiyan Hu, Xiaoman Li

**Author notes:** Corresponding authors. Drs. Haiyan Hu and Xiaoman Li, Harris Corporation and Engineering Center 346, University of Central Florida, Orlando, FL, 32816, USA.

## Abstract

Enhancer-promoter interactions (EPIs) are fundamental to gene regulation, and understanding their recurrence across diverse biological samples is key to deciphering chromatin architecture. In this study, we systematically analyzed the recurrence of EPIs across 49 Hi-C and 95 HiChIP datasets. We found that the majority of EPIs identified in a given sample were also present in other samples, regardless of the assay type (Hi-C or HiChIP) or the enhancer annotations used. Interestingly, EPIs that appeared unique to individual samples were typically surrounded by fewer neighboring EPIs, suggesting they may not represent truly sample-specific interactions. Our findings indicate that most human EPIs have already been captured and that cells primarily reuse subsets of these shared EPIs across different cell types and conditions. This study provides new insights into the pervasive and reusable nature of EPIs in the human genome, with important implications for chromatin conformation studies.

## Introduction

Enhancers are distal regulatory elements that play a pivotal role in controlling gene expression in metazoan genomes^1-3^. Under specific cellular conditions, enhancers can physically interact with their target gene promoters, coming into close spatial proximity of their promoters and significantly boosting the expression of target genes. Given their essential regulatory functions, dysfunction in enhancers or enhancer-promoter interactions (EPIs) have been implicated in various diseases, including cancer and cardiovascular disorders^3-5^. Therefore, understanding the characteristics of EPIs and identifying context-specific EPIs are essential for elucidating gene regulation and disease mechanisms.

To date, many experimental approaches have been developed to map EPIs, primarily based on 3C-derived techniques such as Hi-C, Micro-C, and HiChIP^6-10^. Hi-C and Micro-C can capture a wide range of chromatin interactions, including EPIs, but require deep sequencing to achieve sufficient resolution, making them expensive for large-scale studies. In contrast, promoter-centric approaches such as capture Hi-C and HiChIP are more cost-effective, as they enrich for promoter-associated interactions and require less sequencing to identify EPIs at high resolution^8,9,11^. Nevertheless, the condition-specific and dynamic nature of EPIs, together with their sheer number, makes comprehensive experimental identification across all biological contexts prohibitively costly.

To overcome these limitations, a range of computational methods have been developed for EPI prediction^12-20^. Early models leveraged known features such as conserved synteny and correlated epigenomic signals of enhancer-promoter (EP) pairs^15,18,21-23^. More sophisticated models based on machine learning were later developed to improve the prediction accuracy^14,16,17,24^. Some approaches also infer EPIs directly from chromatin interaction data, though such methods require input experimental data that are often unavailable for cell types and conditions^19,20^.

Despite these advances, the predictive performance of current computational tools for EPI identification remains limited^14,25^. For instance, a recent study showed that even the top-performing methods achieved precision and recall rate of only 0.70 when more than a dozen popular approaches were tested^14^. A key challenge is the vast number of possible EP pairs, which creates a large candidate space that is difficult to filter false EPIs effectively. Prioritizing biologically plausible EP pairs could therefore substantially improve EPI prediction accuracy.

We hypothesize that only a small fraction of EP pairs are truly capable of interacting, and an even smaller subset of these interacting EP pairs is active under specific experimental conditions. This hypothesis stems from the idea that the chemical and physical properties of a DNA sequence impose intrinsic constraints on the number of viable EPIs in a given genomic region. Thus not every EP pair will interact. Even when an EP interaction does occur, it is context-dependent, varying by cell types, tissues, experimental conditions, etc. Current computational methods typically consider all possible EP pairs within a genomic window, which may explain their suboptimal precision and recall^25-27^.

To test this hypothesis, we analyzed EPIs in 95 HiChIP samples and 49 Hi-C samples, the largest datasets we could collect (Material and Methods). We found that EPIs are highly recurrent across samples: on average, over 97.1% of HiChIP EPIs and 88.4% of Hi-C EPIs were observed in multiple samples. Furthermore, for each of the 95 HiChIP samples, at least 77.5% of its EPIs were found in other samples. Similarly, at least 80.4% of the EPIs in each of the 49 Hi-C samples were recurrent. Notably, EPIs detected in only one sample tended to reside in regions with fewer identified EPIs, suggesting that limited sequencing depth in these genomic regions may have hindered their detection in other samples (p-value <0.05). These findings support the notion that the human genome contains a relatively stable set of EPIs, from which individual cell types select subsets for gene regulation. Our study thus provides new insights into the recurrence and reusage of EPIs across samples, with implications for both chromatin architecture and computational EPI prediction.

## Material and Methods

### Collection of chromatin interaction data

We analyzed two types of chromatin interaction datasets. The first dataset consisted of HiChIP loops downloaded from HiChIPDB (https://health.tsinghua.edu.cn/hichipdb/download.php)^28^ (Dated 8/2022, retrieved 10/2024We included all available HiChIP data generated using the H3K27ac antibody, resulting in 95 samples at 5-kilobase resolution. The second dataset comprised Hi-C loops obtained from the 3D genome browser (https://3dgenome.fsm.northwestern.edu/publications.html), which included 49 Hi-C samples at 10-kilobase resolution^29^. We also attempted to obtain chromatin interactions from the 4DN Data Portal (https://data.4dnucleome.org/), but were unable to successfully run its pipelines to generate the required data. As a result, our analysis focused on the 95 HiChIP and 49 Hi-C samples. All chromatin loops from both datasets were mapped to the hg19 genome assembly.

### Enhancers and genes

We used enhancers from two major sources: the Functional Annotation of the Mammalian Genome (FANTOM) project and the Encyclopedia of DNA s (ENCODE)^23,30^. The FANTOM project identified 65,423 enhancers based on bidirectional, capped-RNA transcription of enhancer RNAs, using Cap Analysis of Gene Expression (CAGE) across a wide range of samples^23^. ENCODE computationally predicted 225,753 enhancers based on characteristics of epigenomic signatures in the human genome^30^. Both enhancer sets were annotated with hg19 coordinates. To standardize and better encapsulate enhancer regions^31,32^, we adjusted the lengths of each enhancer to 1000 base pairs if it was shorter than this length^31,32^.

We also obtained 64,864 promoters from GENCODE in the hg19 genome assembly^33^ (Version 46). Each promoter was extended to have a length of 1100 base pairs, with 1000 base pairs upstream and 100 base pairs downstream of the annotated gene transcriptional start site.

### EPIs in 140 cell lines/types

We inferred EPIs from the aforementioned 95 HiChIP samples and 49 Hi-C samples (Supplementary Table S1). We defined EPIs in each of the 144 samples independently. Specifically, for every chromatin loop in a given sample, we identified all enhancers and promoters that overlapped with either loop end. An EP pair was considered to form an EPI in this sample if the enhancer overlapped one end of the loop and the promoter overlapped the other. In this way, multiple EPIs could be defined for a single loop in a sample.

Using FANTOM enhancers, the number of EPIs per sample ranged from 2 to 221,243, with a median of 3,952 and a mean of 24,413. Using ENCODE enhancers, the number of EPIs per sample ranged from 11 to 1,163,985, with a median of 20,177 and a mean of 128,153.

Among these 144 samples, four HiChIP and four Hi-C samples originated from the same cell lines, which were GM12878, K562, KBM7, and THP-1. For these four pairs of shared cell lines, we merged the EPIs from both datasets, removing any duplicates. duplicates This left us with a total of 140 samples for subsequent analysis.

### Repeated occurrence of EPIs across samples

We defined an EPI as shared if it was observed in at least two samples. Conversely, we defined an EPI to be unique if it was observed in only one sample. It is important to note that some EPIs may appear unique due to the limited sequencing depth of certain samples or a small number of available samples.

Since multiple EPIs may be from the same chromatin loop in the above analyses, we employed an alternative method to define shared and unique EPIs based on loop overlap. In this approach, an EPI in one sample was considered shared if its underlying loop was shared by another sample, regardless of whether the same EPI was detected in multiple samples. A loop from one sample was considered shared by a second sample if there existed a loop in the second sample whose two ends each overlapped at least 50% of the corresponding ends of the loop in the first sample. EPIs whose corresponding loops did not meet this overlap criterion were classified as unique.

We further explored the difference between unique and shared EPIs in individual samples. For each sample with at least 100 unique EPIs and 100 shared EPIs, we categorized the EPIs into two groups: unique EPIs and shared EPIs. For each EPI, we counted the number of EPIs that overlapped with it. Two EPIs were considered to overlap if at least one end of one EPI started between the enhancer and the promoter of the other EPI in the genome. We then compared the number of neighboring EPIs for unique and shared EPIs using Mann-Whitney test^34^. If the p-value was less than 0.05, we claimed that shared EPIs had significantly more neighboring EPIs than unique EPIs.

## Results

### An average of ≥97.1% EPIs in each HiChIP sample occurred in other HiChIP samples

We investigated the overlap of EPIs across 95 HiChIP samples, using FANTOM enhancers and GENCODE genes to define the EPIs (Materials and Methods). On average, 97.1% of the EPIs identified in a given sample were also present in other samples, with a median reoccurrence rate of 98.6% (Table 1, Figure 1). In other words, the vast majority of EPIs in any given sample were shared across different HiChIP samples, with recurrency potentially reaching 100%. The sample with the smallest proportion of shared EPIs was COLO320-DM, where 77.5% of its EPIs were found in other samples.

**Table 1:** The summary statistics of the percentages of EPIs that are shared across samples.

| Comparisons | Enhancer types | Minimum | Medium | Maximum | Mean | Standard deviation |
| --- | --- | --- | --- | --- | --- | --- |
| HiChIP | FANTOM | 77.5% | 98.6% | 100.0% | 97.1% | 3.9% |
|  | ENCODE | 86.0% | 98.7% | 100.0% | 97.7% | 2.9% |
| Hi-C | FANTOM | 73% | 81.6% | 100.0% | 82.6% | 5.4% |
|  | ENCODE | 76.8% | 84.1% | 100% | 84.7% | 4.9% |
| Combined | FANTOM | 77.6% | 96.6% | 100.0% | 94.3% | 5.9% |
|  | ENCODE | 81.2% | 97.6% | 100.0% | 95.4% | 5.1% |

Numbers in the first pair of rows were from comparing one HiChIP sample with the remaining 94 HiChIP samples. Numbers in the second pair of rows were from comparing one Hi-C sample with the HiChIP samples that were not from the same cell line. Numbers in the last pair of rows were from comparing one combined sample with the remaining 139 combined samples.

**Figure 1.**
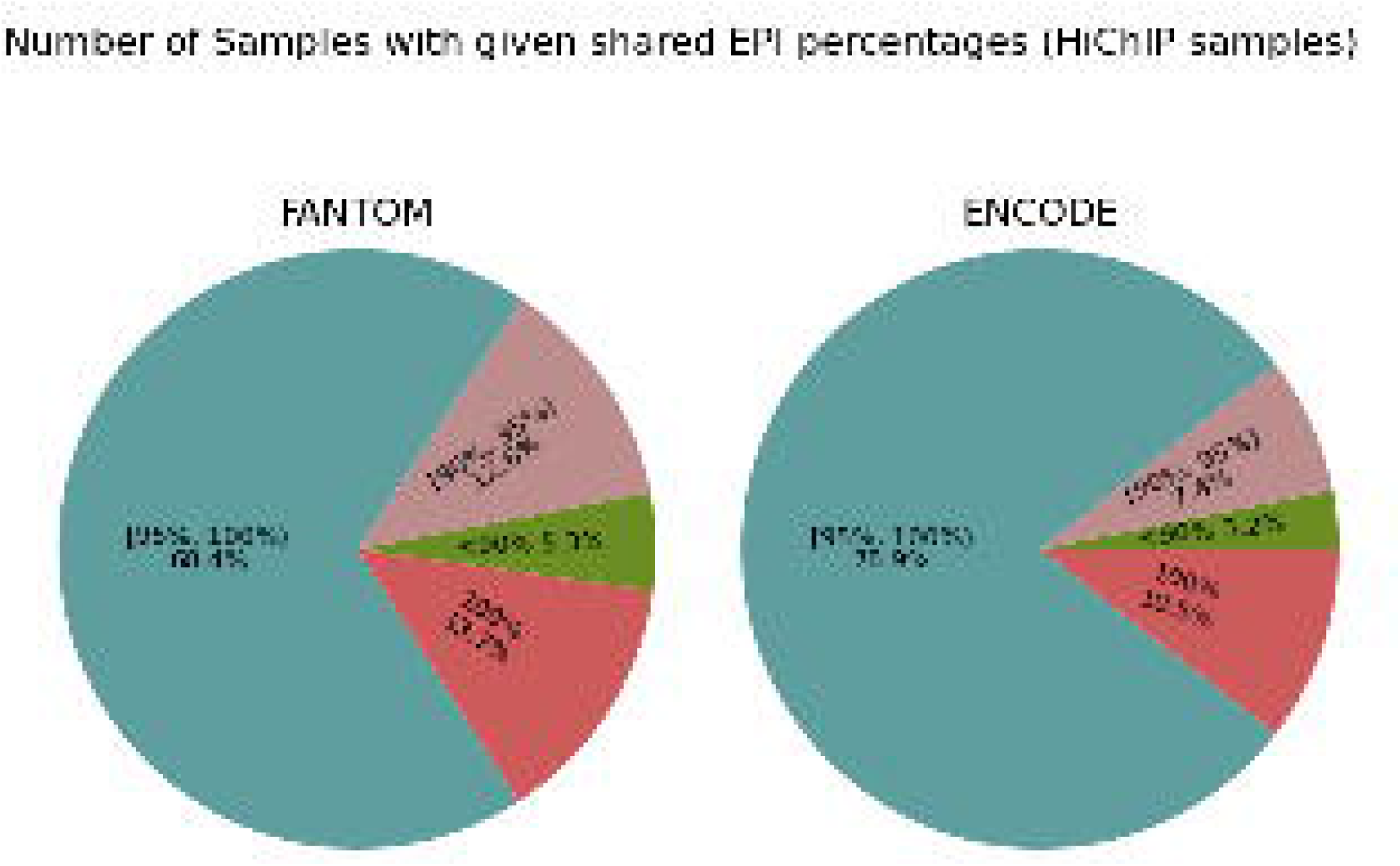
The percentage of EPIs shared in the 95 HiChIP samples. The list of EPIs used in this study are available at https://figshare.com/articles/dataset/EPIs_of_HiChIP_and_Hi-C_samples/29449910

To explore the differences between shared and unique EPIs, we compared the number of neighboring EPIs for these two types of EPIs in every sample that had at least 100 shared and 100 unique EPIs. A total of 51 samples met this criterion. For a given EPI, a neighboring EPI was an EPI with at least one of its ends (promoter or enhancer) overlapping with the genomic region spanned by this given EPI. A Mann-Whitney test revealed a significant difference in the number of neighboring EPIs between shared and unique EPIs in over 96% of the samples (p-value < 0.05). In 79.6% of these samples, shared EPIs had significantly more neighboring EPIs than unique ones (p-value < 0.05), likely due to insufficient sequencing depth in these samples. In contrast, in the remaining 20.4% of samples, shared EPIs had significantly fewer neighboring EPIs than unique ones (p-value < 0.05). Interestingly, these 20.4% of samples had a higher sequencing depth (median: 514,136 loops, mean: 535,671.7 loops) than the above 79.6% samples (median: 130503 loops, mean: 295,418.8 loops). This discrepancy suggests that the low sequencing coverage in other samples may have prevented these unique EPIs in the 20.4% samples from being detected in other samples. Overall, the findings indicate that unique EPIs are likely a byproduct of low sequencing coverage, whereas a core set of EPIs consistently regulates local chromatin structures across different cell types.

Next, we investigated how many samples were required to capture all shared EPIs in a given sample. On average, 1.6, 2.1, and 26.4 samples covered 85.0%, 90.0%, and 100.0% of the shared EPIs, respectively. The corresponding median values were 1, 2, and 23, indicating that most EPIs were shared between pairs of samples, reinforcing the idea that cells repeatedly use the same set of EPIs.

Finally, we assessed whether these findings held when using a different set of enhancers. We repeated the analysis using ENCODE-defined enhancers, which are computationally predicted based on chromatin signatures across samples, in contrast to Whereas FANTOM enhancers derived from balanced bidirectional transcripts in CAGE experiments^23^. Using ENCODE enhancers, we identified a larger set of EPIs in the same 95 HiChIP samples. For instance, 133,317 and 196,227 EPIs were identified in GM12878 and K562 using FANTOM enhancers, whereas 607,027 and 1,146,786 EPIs were identified using ENCODE enhancers.

Despite the larger set of EPIs from ENCODE, we observed similar patterns. A median of 98.7% and an average of 97.7% of EPIs in a sample were also present in other samples. In 99.0% of samples with at least 100 unique and shared EPIs (69 samples), unique EPIs had significantly different number of neighboring EPIs from shared ones, with much fewer neighboring EPIs in 89.9% of samples. Additionally, on average, 1.4, 1.8, and 31.0 samples covered 85.0%, 90.0%, and 100.0% of the shared EPIs in a given sample. These results with ENCODE-defined EPIs further support the conclusion that a common set of EPIs is repeatedly used across different cell types, suggesting that the majority of EPIs in any sample are likely already identified.

### ≥83.5% of EPIs in each Hi-C sample occurred in other samples

We extended the above analysis by examining how many EPIs in each of the 49 Hi-C samples were shared with the 95 HiChIP samples. We focused on the recurrence of EPIs in HiChIP samples because HiChIP experiments generally identify more EPIs than Hi-C experiments at the same sequencing depth^9^. Note that we had both Hi-C and HiChIP data for four cell lines. When comparing EPIs from these Hi-C samples with HiChIP samples, we excluded the HiChIP data from the same cell line or cell type to avoid bias.

Using FANTOM enhancers and GENCODE genes to define EPIs (Materials and Methods), we found that a median of 70.4% and a mean of 71.0% of EPIs in one Hi-C sample were present in other HiChIP samples. When using ENCODE enhancers and GENCODE genes, these values increased slightly, with a median of 72.7% and a mean of 72.9% of EPIs in a Hi-C sample were present in other HiChIP samples. Despite the inherent differences between Hi-C and HiChIP experiments, these results suggest that the majority of EPIs in a Hi-C sample are present in other samples.

The lower recurrence of Hi-C EPIs in HiChIP samples is likely due to the lower resolution of Hi-C data and the imperfect nature of Hi-C-defined EPIs. Both HiChIP and Hi-C experiments primarily define chromatin loops rather than EPIs. Given the lower resolution of the Hi-C data, each loop may overlap with multiple enhancers and/or promoters, leading to the identification of multiple EPIs for a single loop. However, the interaction between two genomic regions represented by a loop may correspond to a single EPI rather than multiple distinct ones. To alleviate this issue of defining EPIs, we revisited the above analyses by considering an EPI in one sample as shared with another sample if the same loop appeared in both samples, even if the EPIs were not identical. Two loops were considered “the same” if more than half of their respective ends overlapped. With this adjusted criterion, we found that, using FANTOM enhancers, a median of 81.6% and a mean of 82.6% of EPIs in a Hi-C sample were present in other HiChIP samples. For ENCODE enhancers, these values were 84.1% and 84.6%, respectively. These results suggest that at least 82.6% of EPIs were shared (Table 1).

Next, we compared shared versus unique EPIs in individual Hi-C samples. We found that unique EPIs had significantly different number of neighboring EPIs from shared EPIs in 90.0% and 100.0% of the samples when using FANTOM and ENCODE enhancers, respectively, where unique EPIs had significantly fewer neighboring EPIs than shared EPIs in 81.8% and 87.5% of samples, respectively. This finding mirrors our previous observation with HiChIP data, suggesting that unique EPIs are likely the result of insufficient sequencing depth in the corresponding regions.

Finally, we examined how many HiChIP samples were required to capture 85.0%, 90.0%, and 100.0% of the shared EPIs in the 49 Hi-C samples. Using FANTOM enhancers, on average, 3.8, 5.4, and 27.4 HiChIP samples captured more than 85.0%, 90.0%, and 100.0% of the shared EPIs in a Hi-C sample, respectively. With ENCODE enhancers, the corresponding values were 3.6, 5.3, and 32.3 HiChIP samples. These relatively small number of required samples suggest that the number of EPIs in a genome is limited, reinforcing the idea that we have already identified most of the EPIs in the human genome. However, it remains unclear which subset of these EPIs is active in a given cell type.

### ≥88.4% of EPIs in a sample were observed in other samples

In previous sections, we explored the recurrence of EPIs within HiChIP or Hi-C samples using different enhancer sets. Since HiChIP experiments generally identify more EPIs than Hi-C experiments at the same sequencing depth, but Hi-C utilizes a distinct technique to capture chromatin interactions^9^, we were interested in worth examining how the combined set of EPIs from both types of experiments affects the recurrence of EPIs across samples. We considered the merged EPIs from 140 samples, including four that contained data from both Hi-C experiments and HiChIP experiments (Material and Methods).

We first compared EPIs identified in Hi-C and HiChIP samples. Among the 95 HiChIP and 49 Hi-C samples, four were common: GM12878, K562, KBM7, and THP-1 (Table 2). In these shared samples, we observed that HiChIP identified at least an order of magnitude more EPIs than Hi-C, and this pattern held for both FANTOM and ENCODE enhancers. For instance, the ratio of HiChIP EPIs to Hi-C EPIs was 34.4 with FANTOM enhancers and 30.9 with ENCODE enhancers. Furthermore, we found that at least 31.0% of EPIs detected in Hi-C experiments were not captured by HiChIP, highlighting that the two techniques identify different subsets of EPIs. This emphasizes the value of integrating data from both methods for a more comprehensive analysis of EPI recurrence across samples. Interestingly, THP-1 showed a much lower percentage of shared EPIs, likely due to its lower sequencing depth and fewer identified EPIs compared with other samples.

**Table 2.** Comparison of EPIs from Hi-C and HiChIP experiments in four samples.

| Samples | Enhancer type | #HiChIP EPIs | #Hi-C EPIs | #shared EPIs | % (#shared/ #Hi-C EPIs) |
| --- | --- | --- | --- | --- | --- |
| GM12878 | FANTOM | 157656 | 4589 | 3156 | 68.8 |
|  | ENCODE | 712107 | 23047 | 14333 | 62.2 |
| K562 | FANTOM | 231825 | 4127 | 2503 | 60.6 |
|  | ENCODE | 1370259 | 22618 | 15555 | 68.8 |
| KBM7 | FANTOM | 166384 | 3787 | 2142 | 56.6 |
|  | ENCODE | 843661 | 20619 | 11156 | 54.1 |
| THP-1 | FANTOM | 66891 | 6143 | 2996 | 48.8 |
|  | ENCODE | 288334 | 30374 | 11724 | 38.6 |

Next, we examined the recurrence of EPIs across the remaining 139 samples (Table 1). We found that a median of 96.6% and a mean of 94.3% of EPIs in a sample were present in other samples when using FANTOM enhancers. With ENCODE enhancers, the median was 97.6% and the mean was 95.4%. For the 49 Hi-C samples alone, the percentage of recurring EPIs was 88.4% for FANTOM enhancers and 88.9% for ENCODE enhancers. These higher recurrence rates, compared to those observed in the second Results section, suggest that Hi-C and HiChIP EPIs complement each other. This further supports the idea that the majority of EPIs in the human genome may have already been identified in published datasets.

Finally, we revisited the comparison between shared and unique EPIs in individual samples. In over 98.0% of the samples with at least 100 unique and shared EPIs, unique EPIs had significantly different number of neighboring EPIs from shared EPIs, whether using FANTOM or ENCODE enhancers. In 88% of these samples, unique EPIs had significantly fewer neighboring EPIs than shared EPIs. This pattern mirrors the previous findings for EPIs derived from individual techniques. Together, these results suggest that we are close to identifying all EPIs in the human genome, although we still do not know which specific subset of these EPIs is active in any given cell type or condition.

## Discussion

In this study, we investigated the recurrence of EPIs across 49 Hi-C and 95 HiChIP samples, considering two different enhancer definitions. We consistently observed that almost all EPIs identified in one sample also appeared in other samples, whether using Hi-C, HiChIP, or the combined data. This result held true regardless of whether FANTOM or ENCODE enhancers were used to define EPIs. Notably, EPIs that appeared unique to individual samples either had significantly fewer neighboring EPIs than those shared across multiple samples, or were detected only in samples with substantially higher sequencing depth and were absent in others with lower coverage. This pattern suggests that truly sample-specific EPIs may not exist; instead, many of these “unique” EPIs could become detectable with deeper sequencing in their genomic neighborhoods. Taken together, our results indicate that the vast majority of human EPIs may have already been identified.

Our findings carry several important implications. First, the number of EPIs in the human genome appears finite. For instance, using FANTOM enhancers, we identified 609,838 EPIs across the 140 combined samples, accounting for only 0.2% of all possible EP pairs within 2 megabase of the genome. Importantly, these EPIs already captured more than 88.4% of EPIs in each individual sample. Second, cells seem to reuse subsets of these shared EPI repertoires. On average, only two samples were sufficient to recover 85.0% of the EPIs found in any given sample, underscoring the extensive overlap among cell types and conditions. Third, because most EPIs are already present in public datasets, computational prediction can now focus on modeling their recurrence across contexts rather than enumerating all possible pairs. Restricting predictions to the observed EPI set eliminates a large number of unlikely interactions, which should substantially improve accuracy in complex mammalian genomes.

Looking ahead, the growing availability of high-throughput chromatin interaction datasets will provide stronger tests of our hypothesis. While we attempted to incorporate additional data, including those from the 4DN portal^35^, we were unable to access relevant datasets at this stage. Incorporating corresponding epigenomic data would further improve EPI definition and resolution^16,17,36-38^. Currently, the lack of matched epigenomic profiles limited the number of samples we could analyze, but this barrier should diminish as more such data become available. With richer epigenomic integration, it will be possible to pinpoint the precise EPIs underlying chromatin loops and conduct more accurate cross-sample comparisons. Finally, although our study supports the view that the repertoire of potential EPIs is limited, the challenge now lies in identifying which subset is active in specific cell types, states, and conditions. Progress in deep learning and other advanced computational methods holds great promise for addressing this challenge, and we anticipate that the next generation of EPI prediction tools will yield more precise and context-aware insights^39,40^.

## Data Availability

The data that supports the findings of this study are available from the following links: Hi-C chromatin loops:

https://3dgenome.fsm.northwestern.edu/downloads/loops-hg19.zip

HiChIP chromatin loops:

https://health.tsinghua.edu.cn/hichipdb/download/fithichip_5k/ChIP/H3K27ac/loop_info.tar.gz

FANTOM5 Enhancers: https://fantom.gsc.riken.jp/5/datafiles/latest/extra/Enhancers/

ENCODE Enhancers:

http://screen.encodeproject.org/

GENCODE Promoters:

https://ftp.ebi.ac.uk/pub/databases/gencode/Gencode_human/release_46/GRCh37_mapping/gencode.v46lift37.basic.annotation.gtf.gz

## Author Contributions

Conceptualization: HH and XL. Formal analysis and investigation: MW, SG, HH, and XL. Resources: HH and XL. Writing-original draft preparation: MW, HH, and XL. Writing-review and editing: MW, HH, and XL. Supervision: HH and XL. All authors read and approved the manuscript.

## Additional Information

## References

1 Howard, M. L. & Davidson, E. H. cis-Regulatory control circuits in development. Developmental biology 271, 109–118, doi:10.1016/j.ydbio.2004.03.031 (2004).

2 Davidson, E. The Regulatory Genome: Gene Regulatory Networks in Development and Evolution. 1 edn, (MA: Academic Press, 2006).

3 Pennacchio, L. A., Bickmore, W., Dean, A., Nobrega, M. A. & Bejerano, G. Enhancers: five essential questions. Nature reviews. Genetics 14, 288–295, doi:10.1038/nrg3458 (2013).

4 Corradin, O. & Scacheri, P. C. Enhancer variants: evaluating functions in common disease. Genome medicine 6, 85, doi:10.1186/s13073-014-0085-3 (2014).

5 Maurano, M. T. et al. Systematic localization of common disease-associated variation in regulatory DNA. Science 337, 1190–1195, doi:10.1126/science.1222794 (2012).

6 Dekker, J., Rippe, K., Dekker, M. & Kleckner, N. Capturing chromosome conformation. Science 295, 1306–1311, doi:10.1126/science.1067799 (2002).

7 Dixon, J. R. et al. Topological domains in mammalian genomes identified by analysis of chromatin interactions. Nature 485, 376–380, doi:10.1038/nature11082 (2012).

8 Javierre, B. M. et al. Lineage-Specific Genome Architecture Links Enhancers and Non-coding Disease Variants to Target Gene Promoters. Cell 167, 1369–1384 e1319, doi:10.1016/j.cell.2016.09.037 (2016).

9 Mumbach, M. R. et al. HiChIP: efficient and sensitive analysis of protein-directed genome architecture. Nature methods 13, 919–922, doi:10.1038/nmeth.3999 (2016).

10 Hsieh, T. H. et al. Mapping Nucleosome Resolution Chromosome Folding in Yeast by Micro-C. Cell 162, 108–119, doi:10.1016/j.cell.2015.05.048 (2015).

11 Li, X., Zheng, Y., Hu, H. & Li, X. Integrative analyses shed new light on human ribosomal protein gene regulation. Scientific reports 6, 28619, doi:10.1038/srep28619 (2016).

12 Roy, S. et al. A predictive modeling approach for cell line-specific long-range regulatory interactions. Nucleic acids research 43, 8694–8712, doi:10.1093/nar/gkv865 (2015).

13 Cao, Q. et al. Reconstruction of enhancer-target networks in 935 samples of human primary cells, tissues and cell lines. Nature genetics 49, 1428–1436, doi:10.1038/ng.3950 (2017).

14 Gschwind, A. R. et al. An encyclopedia of enhancer-gene regulatory interactions in the human genome. bioRxiv, 2023.2011.2009.563812, doi:10.1101/2023.11.09.563812 (2023).

15 Rodelsperger, C. et al. Integrative analysis of genomic, functional and protein interaction data predicts long-range enhancer-target gene interactions. Nucleic acids research 39, 2492–2502, doi:10.1093/nar/gkq1081 (2011).

16 Talukder, A., Saadat, S., Li, X. & Hu, H. EPIP: a novel approach for condition-specific enhancer–promoter interaction prediction Bioinformatics 35, 3877–3883 (2019).

17 Whalen, S., Truty, R. M. & Pollard, K. S. Enhancer-promoter interactions are encoded by complex genomic signatures on looping chromatin. Nature genetics 48, 488–496, doi:10.1038/ng.3539 (2016).

18 Zhao, C., Li, X. & Hu, H. PETModule: a motif module based approach for enhancer target gene prediction. Scientific reports 6, 30043, doi:10.1038/srep30043 (2016).

19 Lu, L. et al. Robust Hi-C Maps of Enhancer-Promoter Interactions Reveal the Function of Non-coding Genome in Neural Development and Diseases. Molecular cell 79, 521–534 e515, doi:10.1016/j.molcel.2020.06.007 (2020).

20 Ron, G., Globerson, Y., Moran, D. & Kaplan, T. Promoter-enhancer interactions identified from Hi-C data using probabilistic models and hierarchical topological domains. Nature communications 8, 2237, doi:10.1038/s41467-017-02386-3 (2017).

21 Thurman, R. E. et al. The accessible chromatin landscape of the human genome. Nature 489, 75–82, doi:10.1038/nature11232 (2012).

22 Malin, J., Aniba, M. R. & Hannenhalli, S. Enhancer networks revealed by correlated DNAse hypersensitivity states of enhancers. Nucleic acids research 41, 6828–6838, doi:10.1093/nar/gkt374 (2013).

23 Andersson, R. et al. An atlas of active enhancers across human cell types and tissues. Nature 507, 455–461, doi:10.1038/nature12787 (2014).

24 Wall, B. P. G., Nguyen, M., Harrell, J. C. & Dozmorov, M. G. Machine and Deep Learning Methods for Predicting 3D Genome Organization. Methods Mol Biol 2856, 357–400, doi:10.1007/978-1-0716-4136-1_22 (2025).

25 Cao, F. & Fullwood, M. J. Inflated performance measures in enhancer-promoter interaction-prediction methods. Nature genetics 51, 1196–1198, doi:10.1038/s41588-019-0434-7 (2019).

26 Xi, W. & Beer, M. A. Local epigenomic state cannot discriminate interacting and non-interacting enhancer-promoter pairs with high accuracy. PLoS computational biology 14, e1006625, doi:10.1371/journal.pcbi.1006625 (2018).

27 Wang, S., Hu, H. & Li, X. Shared distal regulatory regions may contribute to the coordinated expression of human ribosomal protein genes. Genomics 112, 2886–2893, doi:10.1016/j.ygeno.2020.03.028 (2020).

28 Zeng, W. W., Liu, Q., Yin, Q. J., Jiang, R. & Wong, W. H. HiChIPdb: a comprehensive database of HiChIP regulatory interactions. Nucleic acids research 51, D159–D166, doi:10.1093/nar/gkac859 (2023).

29 Wang, Y. L. et al. The 3D Genome Browser: a web-based browser for visualizing 3D genome organization and long-range chromatin interactions. Genome biology 19, doi:10.1186/S13059-018-1519-9 (2018).

30 Consortium, E. P. et al. Expanded encyclopaedias of DNA elements in the human and mouse genomes. Nature 583, 699–710, doi:10.1038/s41586-020-2493-4 (2020).

31 Blanchette, M. et al. Genome-wide computational prediction of transcriptional regulatory modules reveals new insights into human gene expression. Genome research 16, 656–668, doi:10.1101/gr.4866006 (2006).

32 Cai, X. et al. Systematic identification of conserved motif modules in the human genome. BMC genomics 11, 567, doi:10.1186/1471-2164-11-567 (2010).

33 Frankish, A. et al. GENCODE reference annotation for the human and mouse genomes. Nucleic acids research 47, D766–D773, doi:10.1093/nar/gky955 (2019).

34 Mann, H. B. & Whitney, D. R. On a Test of Whether one of Two Random Variables is Stochastically Larger than the Other. Annals of Mathematical Statistics 18, 50–60, doi:10.1214/aoms/1177730491 (1947).

35 Dekker, J. et al. Spatial and temporal organization of the genome: Current state and future aims of the 4D nucleome project. Molecular cell 83, 2624–2640, doi:10.1016/j.molcel.2023.06.018 (2023).

36 Meuleman, W. et al. Index and biological spectrum of human DNase I hypersensitive sites. Nature 584, 244-+, doi:10.1038/s41586-020-2559-3 (2020).

37 Zheng, Y., Li, X. & Hu, H. Comprehensive discovery of DNA motifs in 349 human cells and tissues reveals new features of motifs. Nucleic acids research 43, 74–83, doi:10.1093/nar/gku1261 (2015).

38 Wang, Y., Li, X. & Hu, H. H3K4me2 reliably defines transcription factor binding regions in different cells. Genomics 103, 222–228, doi:10.1016/j.ygeno.2014.02.002 (2014).

39 Athaya, T., Ripan, R. C., Li, X. M. & Hu, H. Y. Multimodal deep learning approaches for single-cell multi-omics data integration. Briefings in bioinformatics 24, doi:10.1093/bib/bbad313 (2023).

40 Talukder, A., Barham, C., Li, X. M. & Hu, H. Y. Interpretation of deep learning in genomics and epigenomics. Briefings in bioinformatics 22, doi:10.1093/bib/bbaa177 (2021).

