## Supplementary Table S1 for "Recurrent enhancer-promoter interactions across samples"

| Enhancer | Samples | min | median | max | mean | std dev |
| --- | --- | --- | --- | --- | --- | --- |
| FANTOM | Hi-C only | 18 | 2218 | 8279 | 2672.92 | 1602.63 |
| ENCODE |  | 96 | 9153 | 38802 | 12557.94 | 8465.29 |
| FANTOM | HiChIP only | 2 | 10108 | 221243 | 34670.36 | 54261.2 |
| ENCODE |  | 11 | 60676 | 1163985 | 182745.41 | 283096.44 |
| FANTOM | both | 2 | 3952 | 221243 | 24412.64 | 47258.9 |
| ENCODE |  | 11 | 20177 | 1163985 | 128152.76 | 247063.67 |

supplementary table S1: Statistic Summary of the number of EPIs found within the 140 samples collected from both Hi-C and HiChIP (Duplicate samples are combined).
